# Buyer Beware: confounding factors and biases abound when predicting omics-based biomarkers from histological images

**DOI:** 10.1101/2024.06.23.600257

**Authors:** Muhammad Dawood, Kim Branson, Sabine Tejpar, Nasir Rajpoot, Fayyaz ul Amir Afsar Minhas

## Abstract

**Background:** Recent advancements in computational pathology have introduced deep learning methods to predict genomic, transcriptomic and molecular biomarkers from routine histology whole slide images (WSIs) for cancer diagnosis, prognosis, and treatment. However, existing methods often overlook the critical role of co-dependencies among biomarker statuses during training and inference. We hypothesize that this oversight results in models that predict the combined effect of multiple interdependent biomarkers rather than individual statuses independently, akin to attributing the quality of an orchestral symphony to a single instrument, highlighting limitations of current predictors.

**Methods:** Using large datasets (n = 8,221 patients), we conducted statistical co-dependence testing to demonstrate significant interdependencies among biomarker statuses in training datasets. Following standard protocols, we trained two machine learning models to predict biomarkers from WSIs achieving or matching state-of-the-art predictive performance. We then employed permutation testing and stratification analysis to evaluate their predictive quality based on the principle of conditional independence, i.e., if a model accurately captures the phenotypic influence of a specific biomarker independent of other biomarkers, its performance should remain consistent across subgroups of patients stratified by other biomarkers, aligning with its overall performance on the entire dataset.

**Findings:** Our statistical analysis reveals significant interdependencies among biomarkers, reflecting expected co-occurrence and mutual exclusivity patterns influenced by pathological and biological processes that are consistent across datasets, as well as sampling artefacts that can be different across datasets. Our results indicate that the predictive quality of an image-based predictor for a biomarker is contingent on the status of other biomarkers, revealing that models capture aggregated influences rather than predicting individual statuses independently. For example, mutation predictions are confounded by the overall tumour mutation burden. We also show that, due to the presence of such correlations, deep learning models may not offer significant advantages in predicting certain biomarkers in comparison to simply using pathologist-assigned grades for their prediction.

**Interpretation:** We show that current deep learning models in computational pathology fall short in isolating individual biomarker effects, leading to confounded and less precise predictions. Our findings suggest revisiting model training protocols to recognize and adjust for biomarker interdependencies at all development stages—from problem definition to usage guidelines. This involves selecting diverse datasets to reflect clinical heterogeneity, defining prediction variables or grouping patients based on co-dependencies, designing models to disentangle complex relationships, and stringent stratification testing. Clinically, failure to account for interdependencies may lead to suboptimal decisions, necessitating appropriate usage guidelines for predictive models.

## 1 Introduction

In pace with developments in computational pathology, several studies have emerged in recent years aiming to predict clinically relevant biomarkers^1–5^ such as gene mutations and expression levels from histology images^1–9^. These approaches take a whole-slide image (WSI) as input and predict the status of clinically relevant biomarkers such as microsatellite instability (MSI), estrogen receptor (ER) and progesterone receptor (PR) statuses or the presence of alterations in individual genes such as TP53, BRAF, KRAS, EGFR etc as their target. Typically, the goal of these predictors is to predict the status of molecular biomarkers and/or to identify or mine histological structures or patterns associated with those biomarkers^10^. Another clinically driven motivation of such methods is to rule out certain gene mutations using routine histology images without the need for additional stains or molecular testing, both of which can be more costly, can be tissue destructive and present significantly longer waiting times for patients^11^. For example, accurate prediction of MSI^1,5,12–14^ and mutations in BRAF^1^ and KRAS/NRAS^6^ genes from histology images can aid in tailoring clinical decisions for personalised therapeutic intervention, thereby reducing the cost and waiting time compared to sequencing^11^.

Several methods have shown that it is possible, at least in a statistically significant sense, to predict biomarkers status^2,3,15^ and somatic alterations in numerous genes from WSIs in specific cancers^6–8,16^. Most methods in this domain utilise deep learning pipelines with weakly supervised learning for training a predictor on WSIs from The Cancer Genome Atlas (TCGA) or other similar data repositories such as the Clinical Proteomic Tumour Analysis Consortium (CPTAC)^17^. The predictive accuracy of a majority of current methods for most clinically relevant biomarkers and gene mutations currently remains rather low with reported performance metrics such as the Area Under the Receiver Operating Characteristic curve (AUROC) in the range of 0.50-0.90. When technical factors such as mutation prevalence, scarcity of multi-centric data, the class imbalance between positive (mutated or highly expressed) and negative (wild-type or low expression) cases, quality of images (pen markings, tissue tears, etc), robustness, domain shifts across different datasets, and generalisation to unseen datasets from other centres are considered, the true generalisation of such histology image-based predictors to large unseen datasets faces significant challenges. In this paper, we demonstrate that even if these challenges have been handled, there are underlying fundamental issues that require addressing. Specifically, we highlight the importance of considering interdependence between the status of different omics-based biomarkers in terms of their co-occurrence and mutual exclusivity when predicting their status from WSIs through machine learning.

There are two fundamental reasons for the emergence of such interdependencies or associations among different biomarkers or mutations: 1) these can be causal or persistent, i.e., stem naturally from genomic or transcriptomic pathways or mutations underlying disease progression, e.g., the co-occurrence of certain gene mutations in tumours or 2) be spurious, confounding or non-causal due to limited or biased samples or due to differences between training and test data. It can be very difficult to discern between these two types of associations purely from observational data. However, such interdependencies manifest themselves as correlation or mutual information between these factors. For example, in most solid tumours (e.g., breast, colon), around 33 to 66 genes undergo inter-dependent somatic alterations, which are likely to alter their protein expression^18^. Tumours typically accumulate these mutations progressively over time when they evolve from a benign lesion to a malignant tumour^19,20^. This process is well studied in colorectal cancer where an initial mutation in the APC gene provides a selective growth advantage to epithelial cells, leading to the formation of adenoma^21^. Subsequently, mutations in other genes (e.g., KRAS, PIK3CA, SMAD4 and TP53) further accelerate tumour growth, eventually resulting in a malignant tumour capable of invasion and metastasis. A similar pattern has also been observed in breast tumours, where higher-grade tumours generally harbour a higher number of mutations compared to low-grade tumours^22^. The co-occurrence of multiple somatic mutations in tumours reflects the concept of *epistasis* where gene interactions produce a synergistic effect on the phenotype^23–26^. Tumour evolution favours random mutations that offer fitness advantages. Mutations in genes with *positive epistasis* result in strong fitness compared to their individual fitness effects and are more likely to occur together in the same tumour (co-occurrence). In contrast, mutations in genes with *negative epistasis* yield a surprisingly weak fitness compared to their individual effect and are rarely co-present within the same tumour resulting in mutual exclusivity. For example, in lung cancer TP53, RB1 and EGFR^27^ are often co-mutated, whereas mutations in KRAS and EGFR show a pattern of mutual exclusivity^24,25,28^. Such co-dependencies and multifactor drivers of phenotypes are unavoidable. However, these patterns can also differ from one study or dataset to another depending on sample size, patient selection, treatment regimens etc. It can be quite difficult to pin a phenotypic effect in a histology image to a single molecular factor. Consequently, such confounding or non-causal differences can be a source of generalization failure of machine learning models that are trained over data in which both causal and non-causal dependencies are present.

In histology images, disease phenotypes emerge from the interaction of multiple co-dependent genes rather than a single factor such as mutation of a single gene. Disentangling the contributions of individual genes to these phenotypes is a complex challenge. Current computational pathology methods that predict the status of gene mutations or gene expression levels from WSIs often focus on single-gene effects and neglect the critical role of gene co-dependencies. This oversight is akin to attributing the quality of a symphony played by an orchestra to a single instrument. Moreover, when the interdependencies among omics-based factors in a test case differ from those in the training data, such a mismatch or domain shift can hinder a machine learning model’s ability to generalise effectively. In this work, we highlight the need for models that can navigate the complex omics-driven landscape of diseases^29^. Our key contributions in this paper are as follows:

We show that complex inter-dependencies between molecular factors exist in real-world data and can be significantly different across datasets indicating the presence of both causal and spurious associations across the molecular factors.

1. Through permutation testing and stratification analysis, we demonstrate failure modes of histology image-based biomarker predictors by showing that their accuracy for predicting the status of certain molecular biomarkers can vary significantly when conditioned on the status of other biomarkers. Reminiscent of the effect of collinearity in statistics, the inter-dependencies among biomarkers in training lead the model to predict the aggregated effect of multiple factors rather than individual ones independently.
2. We recommend making appropriate causal adjustments in histology image-based predictors of molecular status to ensure that correct inferences are made from histology images before making an important decision (such as treatment selection) based on those inferences.

## 2 Results

### 2.1 Analytic workflow

A high-level concept diagram of machine learning approaches for predicting genomic and transcriptomic biomarkers such as mutations, gene expression patterns, genomic instability indicators, etc. of cancer patients from routine histological images and their limitations is provided in **Figure 1**. We argue that the significant interdependence among biomarker statuses across cancer patients and the disregard of such interdependencies in model training makes it challenging for machine learning algorithms to identify individual biomarker-level genotype-to-phenotype mapping independent of other biomarkers. Instead, the predictor is likely to predict the combined effect of multiple biomarkers whose statuses are associated with each other in the training dataset. We demonstrate this with permutation testing and stratification analysis across different cancer types, and cohorts as well as different machine learning methods. The key steps in our analytic workflow are outlined below (see methods section for details).

**Figure 1:**
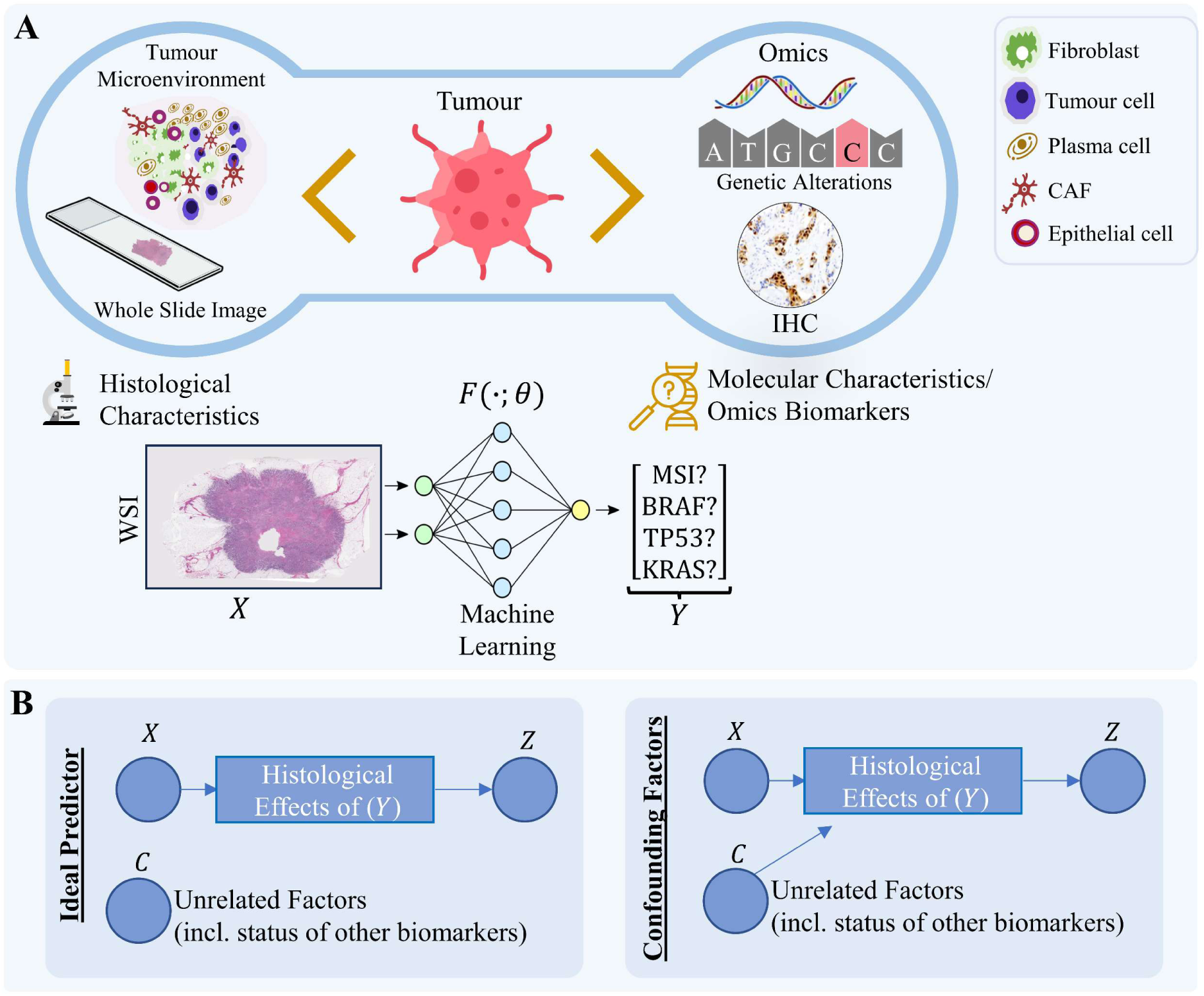
Diagram illustrating the conceptual framework of machine learning methods aimed at predicting the status of molecular biomarkers from histology images. **A)** Machine learning based prediction of molecular characteristics or omics biomarkers from whole slide images involves using training data of WSIs with known biomarker statuses. The machine learning model accepts the representation of the whole slide image (*X*) as input and predicts the status of a certain biomarker (*Y*) as the target. **B)** An ideal predictor should be able to predict mutational or omics-based biomarker signature using features based on the histological effects of that biomarker and its output (*Z*) should be independent of unrelated confounding factors (lumped into a variable *C*) as shown in the simplified causal diagram. Conversely, if the predictor’s output is dependent upon the histological effects of (*Y*) as well as other confounding factors such as histological grade or tumour mutational burden, it may not be possible to tease out the individual effect of Y independently.

We first analysed the interdependency among biomarkers and somatic mutation status of genes in samples (n = 8,221) with breast, colorectal, endometrial and lung cancers across four different cohorts: the cancer genome atlas (TCGA n = 2,683), Molecular Taxonomy of Breast Cancer International Consortium (METABRIC n = 2,433)^30,31^, Memorial Sloan Kettering Cancer Centre (MSK) (n = 2,486)^32,33^ and DFCI (n = 619)^34^. The interdependency analysis assesses co-occurrence and mutual exclusivity associations among biomarkers and gene mutation status within each cohort using log odds ratio (LOR) and Fisher exact test.

Next, we assessed the predictability of biomarkers and gene mutations from WSIs using two different algorithms with different principles of operation: CLAM - which is an attention-based aggregation method^35^ and a graph neural network based approach called *SlideGraph*^∞^ ^36^. These models and their training are representative of the variety of existing machine learning methods in the literature that do not make any explicit design considerations for modelling inter-dependencies between prediction variables. For each biomarker, we evaluate the test performance of both algorithms in the TCGA cohort using AUROC as a performance metric ensuring that these models achieve or match state-of-the-art performance. Additionally, we also assessed the performance of the trained image-based biomarkers and gene mutation predictors on patients in two independent validation cohorts, namely, CPTAC^37^ and the Australian Breast Cancer Tissue Bank (ABCTB)^38^.

Finally, we demonstrate the influence of confounding factors on image-based biomarker predictors across different cancer types and cohorts using stratification analysis and permutation testing. This is based on the principle of conditional independence, i.e., if a model accurately captures the phenotypic influence of a specific biomarker independent of other biomarkers, its performance should remain consistent (within statistical tolerances) across subgroups of patients stratified by other biomarkers, aligning with its overall performance on the entire dataset. Differences between predictive performance within and across the datasets reveal the presence of dependencies between biomarkers showing limitations of predictive models in terms of predicting the aggregated effects of multiple biomarkers. Based on these results, we identify potential failure modes of such models.

### 2.2 Biomarkers and genes mutation status show significant inter-dependency

From our interdependency analysis, we found significant associations (two-sided Fisher exact multiple hypothesis corrected *p* ≪ 0.05) among biomarkers across different cancer subtypes (see **Figure 2** and **Figure S1**). For breast cancer cases in both TCGA breast (TCGA-BRCA) and METABRIC cohorts, we found elevated expression of ER and PR to be co-occurring (high positive log odds ratio) with mutations in CDH1, MAK3K1 and PIK3CA while not with TP53 mutation showing mutually exclusivity. Additionally, TP53 mutation status shows mutual exclusivity with mutations in CDH1, GATA3, MAP3K1 and PIK3CA genes. This observation aligns with a previous study that has found similar patterns of association in breast tumours^39^.

**Figure 2:**
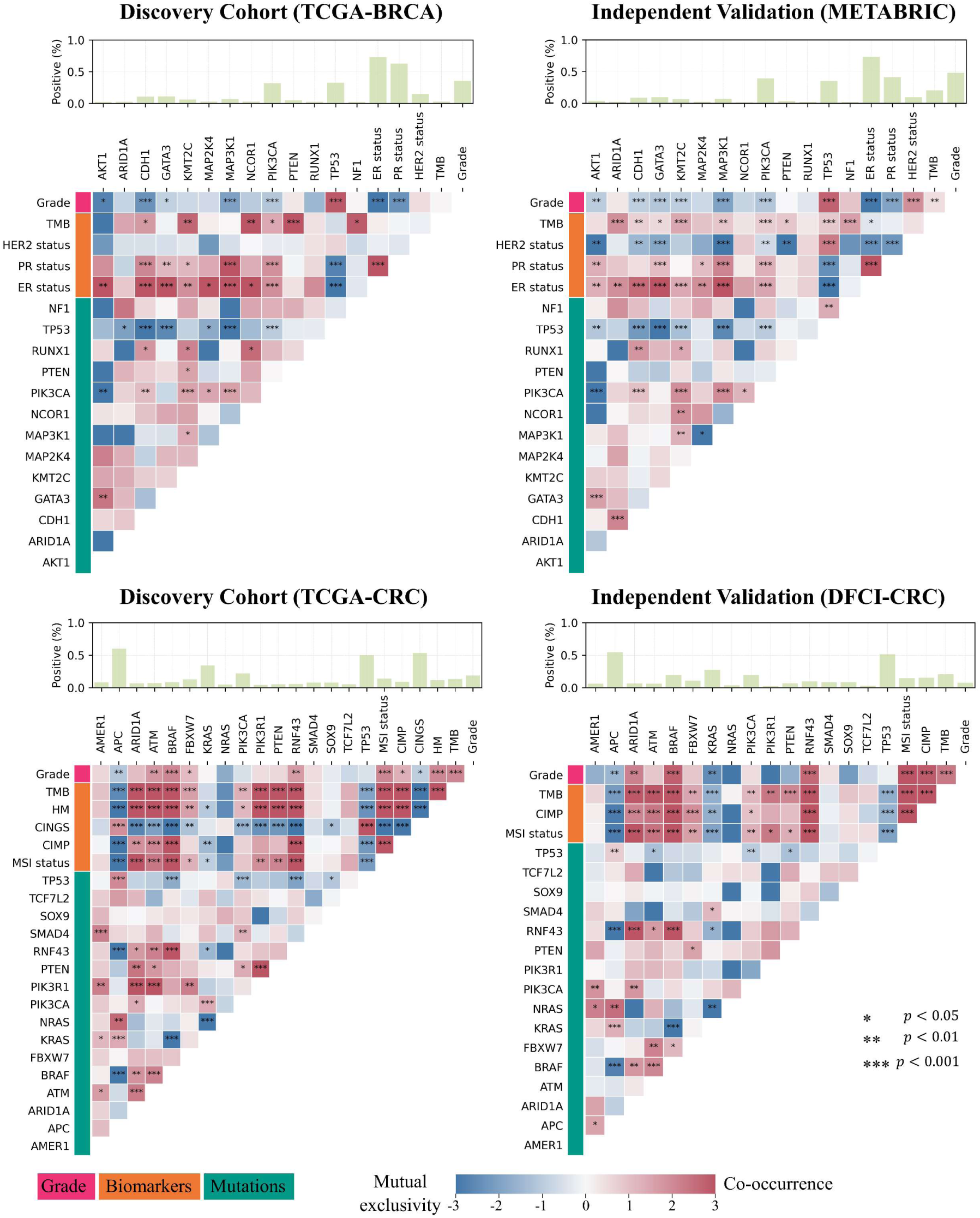
Heatmaps showing association of biomarkers as well as gene mutation statuses across tissue types and datasets. The heatmaps display a set of biomarkers and genes along the axes with cell colours within the heatmap showing the strength of association (dark-red colours for co-occurrence and dark blue for mutual exclusivity). Cells marked with asterisks indicate statistically significant associations (two-sided Fisher’s exact test multiple hypothesis corrected *p* ≪ 0.05). The top bar above each heatmap shows the percentage of cases mutated for a specific gene in case of gene mutations, whereas for biomarkers it indicates the percentage of patients with elevated estrogen receptor (ER), progesterone receptor (PR) and human epidermal growth factor receptor 2 (HER2) in case of breast tumours, high microsatellites instability (MSI), hypermutation and CpG island methylator phenotype pathway (CIMP) activity, and chromosomal instability (CIN) for colorectal tumours. Abbreviations: chromosomal instable vs genome stable (CINGS), hypermutated (HM), tumour mutational burden (TMB).

In colorectal cancer, MSI high cases in both TCGA and DFCI cohorts carry mutations in BRAF, ATM, ARID1A and RNF43 genes, while lacking KRAS mutation (see **Figure 2**). Furthermore, BRAF mutant tumours predominantly exhibit a higher TMB and carry mutations in ATM, RNF43 and ARID1A (see **Figure 2**). Apart from colorectal cancer, we also found significant interdependency (*p* ≪ 0.05) among the mutational status of genes in endometrial cancer and lung adenocarcinoma from the TCGA and MSK cohort (see **Figure S1**). For example, in endometrial cancer, patients with PTEN mutation also carry mutations in APC, ATM, JAK1, KRAS, ARIDA and several other genes. Similarly, lung adenocarcinoma (LUAD) patients with STK11 mutations also have KEAP1 mutation but not EGFR mutation.

This key result lends support to the motivation underlying this work, i.e., biomarkers and genes mutation status show significant interdependency. Consequently, designing an ML model to map phenotypic patterns in WSIs with the status of an individual biomarker or gene mutation status could be challenging, as the observed phenotypic patterns in WSIs might not be exhibited by the status of a singular biomarker or gene mutation but rather it would be a combined effect exhibited by a set of biomarkers of gene mutation that show significant patterns of interdependency. Such co-dependencies in biomarkers can be persistent or causal in which case they are to be expected and inescapable. However, as discussed in the next section, these underlying association structures can vary across datasets and can be sampling artefacts leading to potential confounding effects.

### 2.3 Associations between biomarkers vary across datasets

From our interdependency analysis, we found that the nature of associations among the statuses of different biomarkers can also vary across datasets. For instance, in the TCGA-BRCA cohort MAP3K1 mutation shows mutual exclusivity with AKT1 and ARID1A mutations, whereas in the METABRIC cohort, they show a tendency of co-occurrence (see **Figure 2**). Moreover, elevated ER expression and high TMB show a small degree of co-occurrence in the TCGA-BRCA cohort, whereas, in the METABRIC cohort, they show mutual exclusivity. Similarly, in the TCGA colorectal (TCGA-CRC) cohort, BRAF mutant tumours are significantly (p≪0.05) less likely to exhibit TP53 mutations, however, this association is less pronounced in the DFCI cohort and lacks statistical significance. Additionally, we also found variations in association patterns between biomarkers statuses in endometrial and lung cancer samples across the dataset (see **Figure S1**). For instance, in the TCGA-LUAD cohort, BRAF mutations demonstrate a degree of mutual exclusivity with STK11 mutations, whereas, in the MSK cohort, they exhibit a slight tendency towards co-occurrence.

Such differences in the association patterns among biomarker statuses across datasets may limit ML-models’ generalisability by introducing dataset-specific priors into the model, potentially leading to poor test performance if these patterns vary significantly in the test dataset compared to the training dataset.

### 2.4 Prediction of biomarkers and gene alternations from whole slide images

To analyse the impact of confounding factors on image-based predictors across models and feature types, we trained two different machine learning models (CLAM and *SlideGraph*^∞^) with two different feature representations or embeddings. These models and their training are representative of the variety of existing machine learning methods in the literature in that they do not make any explicit design considerations for modelling inter-dependencies between prediction variables. However, before proceeding with the analysis of confounding factors, it was ensured that these approaches give consistent or higher predictive performance in comparison to similar methods in the literature for different biomarkers and gene mutation predictions. These results are shown in **Figure 3** and **Figure S2**. For instance, CLAM with a self-supervised feature extractor predicts the ER and PR status of patients in cross-validation (TCGA-BRCA) and an independent validation cohort (ABCTB) with average AUROCs of (ER:0.87, PR:0.79) and (ER:0.90, PR:0.78), respectively. Similarly, the same model predicts CDH1 and TP53 mutations status of patients in cross-validation (TCGA-BRCA) and independent validation (CPTAC-BRCA) cohorts with AUROCs of (CDH1:0.88, TP53: 0.82) and (CDH1: 0.91, TP53: 0.82), respectively. In colorectal cancer, *SlideGraph*^∞^with a self-supervised feature extractor, predicts the MSI status of patients in both cross-validation (TCGA-CRC) and independent validation (CPTAC-CRC) cohorts with AUROC values of 0.89 and 0.84, respectively. Furthermore, we also managed to achieve higher AUROC for predicting BRAF mutation, CIMP, CINGS, and HM status from histology images (see **Figure 3**). Besides breast and colorectal cancer, we accurately predicted gene mutations in lung and endometrial cancer samples from histology images with high AUROC values in both cross-validation (TCGA) and independent validation (CPTAC) cohorts (**Figure 3**, **Figure S2**).

**Figure 3:**
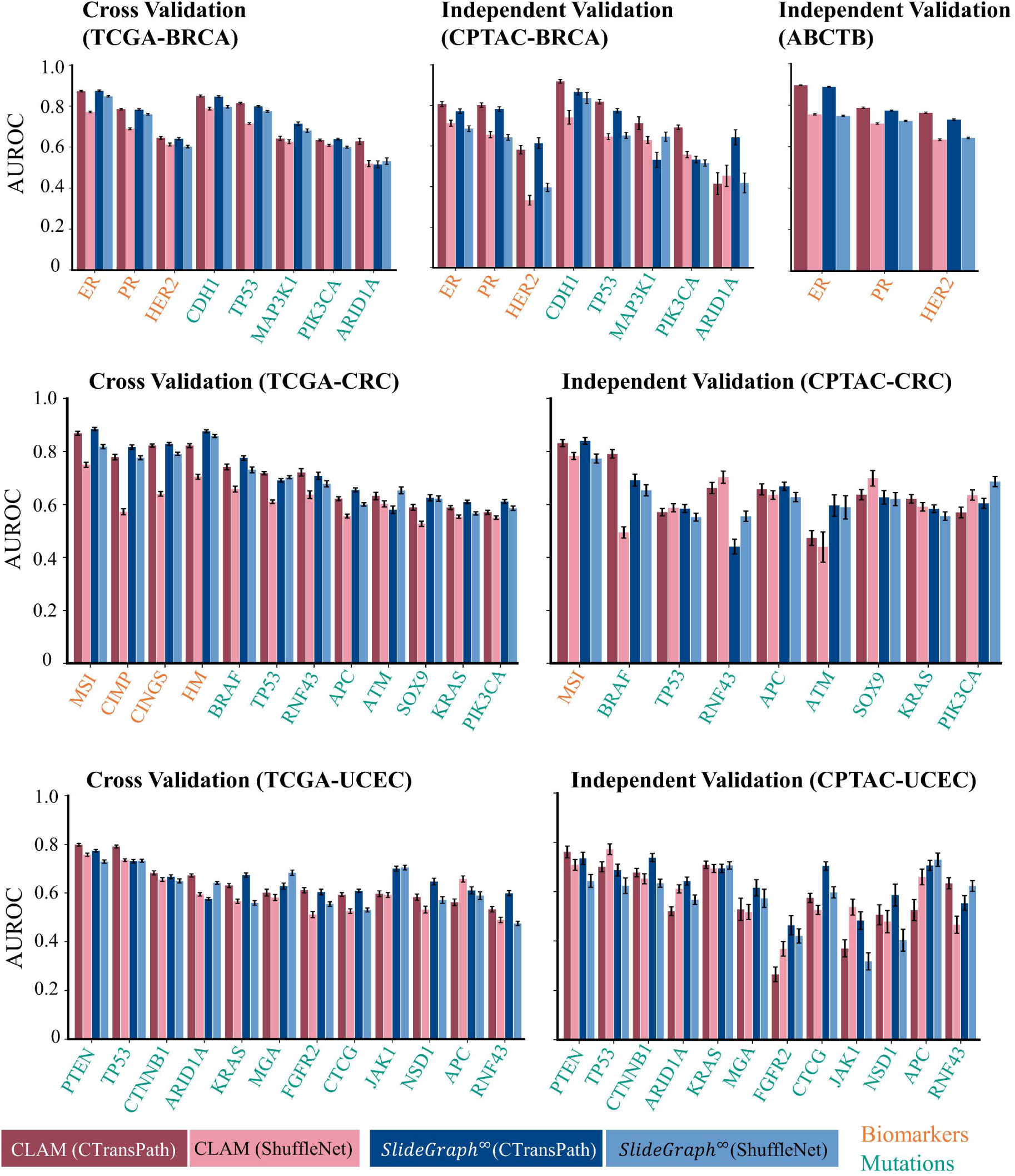
Quantitative results of prediction biomarkers/mutations from histology images. Each barplot shows the area under the receiver operating characteristic curve (AUROC) at which certain clinical markers have been predicted using two weakly supervised models (CLAM and *SlideGraph*^∞^) each with two different feature representations (i.e., using ShuffleNet pretrained on natural images as patch-level encoder, and CTransPath a transformer based model pre-trained on histology images using self-supervised learning).

These results underscore the effectiveness of ML-predictors employed in the study for accurately predicting biomarkers and gene mutations from histology images. Next, we analyse the influence of confounding factors on the prediction of various biomarkers and mutations through these models as illustrative examples of failure modes of similar approaches in computational pathology.

### 2.5 Interdependence in biomarker status leads to entangled histology phenotypes captured from WSIs

Due to the significant inter-dependency among biomarkers and gene mutation status in the training dataset, designing a predictor to predict the phenotypic effect of a singular biomarker or gene mutation from histology images is challenging. This co-dependency is likely to strongly influence histology image-based predictors. To test this hypothesis, we compared the predictive performance of a machine learning model for predicting a certain biomarker (e.g., MSI status in colorectal cancer) over a test cohort with its performance stratified based on statuses of other biomarkers or mutations with significant differences between them indicative of the possibility of confounding or associative factors. For example, while the machine learning model used in this work achieved an AUROC value of 0.88 (0.873-0.886) for predicting colorectal tumours MSI status in colorectal tumours, stratification analysis reveals a significant drop in model predictive performance in certain subgroups. If the patients in the test cohort are divided into hypermutated and non-hypermutated subgroups, the model performance drops from 0.88 across the overall dataset to 0.72 in hypermutated and non-hypermutated cases. A similar effect was observed in stratification by other biomarkers showing co-occurrence—e.g., CPG island methylator phenotype pathway (CIMP) activity, hypermutation (HM) and APC statuses—or mutual exclusivity, e.g., BRAF and chromosomal instable vs genome stable (CINGS) association MSI status (see **Figure 2** and **Figure 4**). This suggests that the predictions of the machine learning model for a given biomarker is dependent upon the status of other biomarkers and that the model may be leveraging the combined phenotypic effect that emerges from potentially interacting biomarkers (**Figure 2**), rather than the one solely linked to MSI status variations.

**Figure 4:**
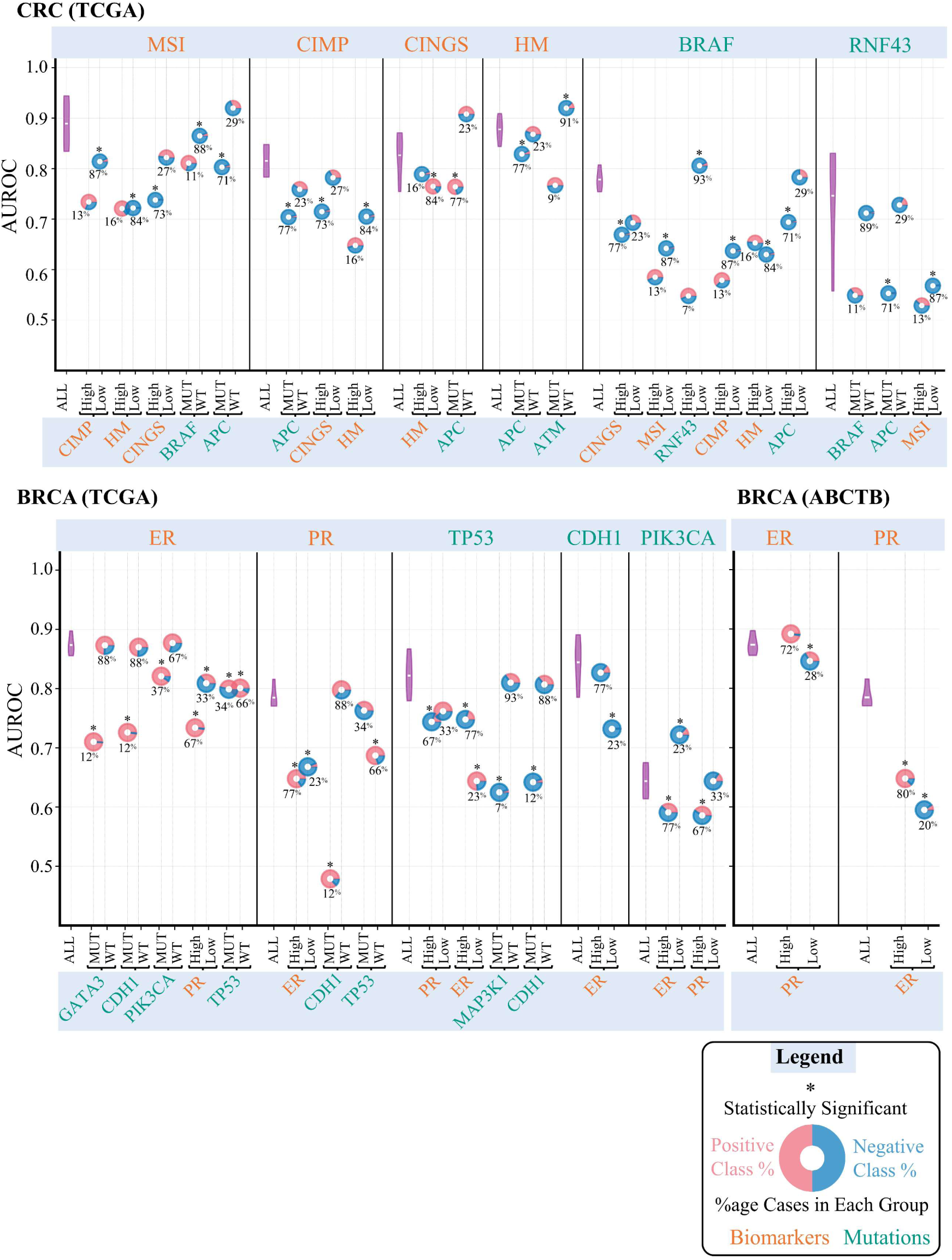
Plots showcasing stratification analysis of several histology image-based biomarker predictors concerning other interdependent biomarkers. In the plots, AUROC values are illustrated on the y-axis, with the top x-axis indicating the prediction variables and the bottom x-axis showing the stratification variables. The predictive performance of each predictor on all the cases in the cohort (denoted by All in the plot) over 100 bootstrap runs is shown using a violin plot, whereas its performance in different stratification groups is depicted with a donut chart, with the centre representing the AUROC values. Donuts marked with * at the top indicate statistically significant results (FDR-corrected p-values of permutation testing *p* ≪ 0.05). The percentage values at the bottom of the donut indicate the proportion of positive (MUT/High) or negative (WT/Low) cases relative to the status of the stratification variables. Red and blue colours in each donut indicate the proportion of positive and negative cases in each stratified group concerning prediction variables.

These observations extend beyond image-based biomarker predictors in colorectal tumours and are also noticeable in biomarker predictors for breast and endometrial cancers, as illustrated in **Figure 4** and **Figure S3**. For instance, in breast tumours, the performance of the ER predictor experiences a notable decline in cases with mutations in GATA3, CDH1, and PIK3CA (see **Figure 4**). Similarly, the ER predictor’s performance diminishes significantly in both PR-positive and negative cases, as well as in cases with and without TP53 mutations. Similar trends are apparent for PR, TP53, CDH1, and PIK3CA predictors, as shown in **Figure 4**.

### 2.6 Breast tumour receptor status predictors are confounded by histology grade

We analysed the influence of histology grade on receptor status predictors by comparing the performance of these models over test cohorts stratified based on patients’ histological grades. Histology image-based models predict the ER and PR statuses of patients in the TCGA-BRCA and ABCTB cohorts with average AUROCs of (ER: 0.87, PR: 0.79) and (ER: 0.90, PR: 0.78), respectively. This indicates image-based models’ strong capability in distinguishing receptor status positive cases from negative cases using routine WSIs. However, these figures, while initially promising, are nuanced by a stratification analysis by tumour grade (see **Figure 5**). For instance, the performance of the ER predictor significantly declines (FDR-corrected p-values of permutation testing p≪0.05) to an AUROC of 0.76 (from 0.87) for medium-grade tumours in both cohorts. Similarly, the PR predictor’s performance significantly decreases in low and medium-grade tumours in both TCGA-BRCA and ABCTB cohorts (see **Figure 5A**). In TCGA-BRCA, its performance reduces to AUROC values of 0.59 and 0.69, while in the ABCTB cohort, the AUROC declines to 0.65 and 0.73, respectively for low and medium-grade tumours. This analysis suggests that the ER model’s predictions in low-grade tumours and the PR model’s predictions in low and medium-grade tumours are not substantially more informative than those derived from manual grade assignment by pathologists (see **Figure 6**).

**Figure 5:**
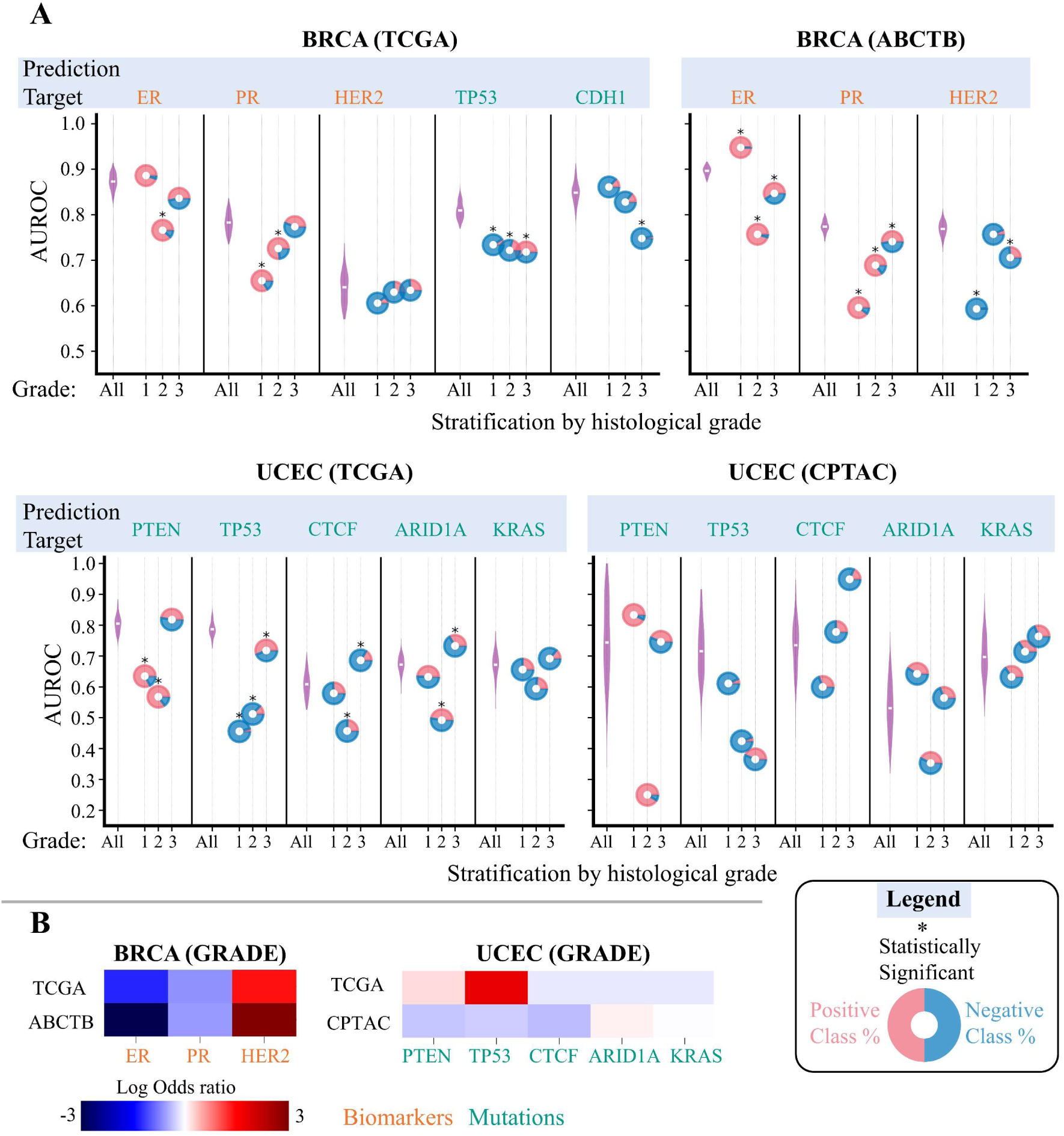
Plots illustrating the biased predictive performance of image-based biomarker predictors across patients with different histological grades through stratification analysis. **A)** In the plots, AUROC values are illustrated on the y-axis, with the top x-axis indicating the prediction variables and the bottom x-axis showing patient stratification with respect to histological grade. The predictive performance of each predictor on all the cases in the cohort (denoted by All in the plot) over 100 bootstrap runs is shown using a violin plot, whereas its performance in a grou-p of patients with a certain histological grade is depicted with a donut chart, with the centre representing the AUROC values. Donuts marked with * at the top indicate statistically significant results (FDR-corrected p-values of permutation testing *p* ≪ 0.05). Red and blue colours in each donut indicate the proportion of positive and negative cases in each stratified group in relation to prediction variables. **B)** Heatmaps highlighting the shift in the association structure between histological grade and biomarkers status across two distinct datasets. Color intensity reflects the strength of association, with dark-red indicating strong co-occurrence and dark-blue indicating strong mutual exclusivity.

**Figure 6:**
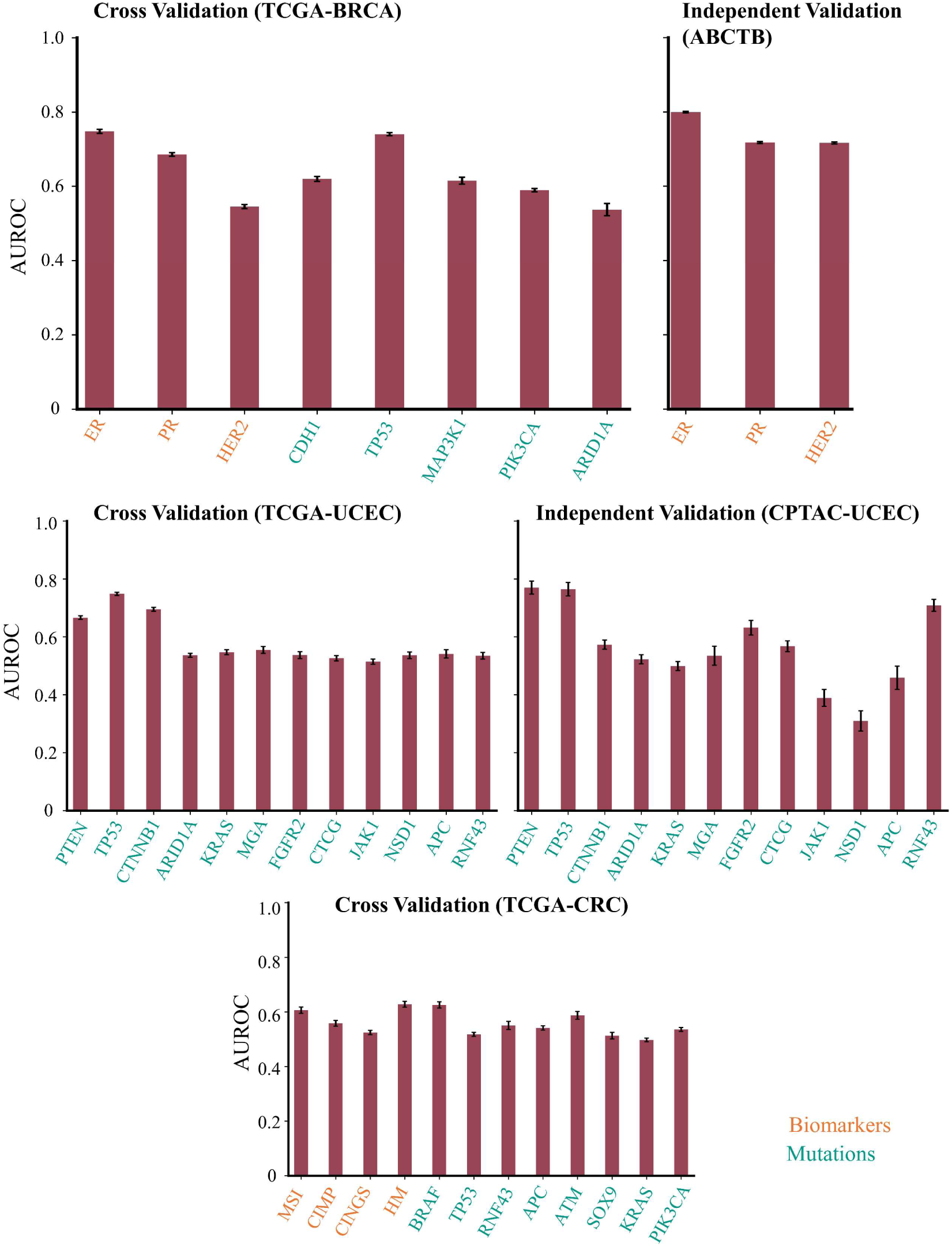
Quantitative results of prediction of biomarkers/mutations using pathologists’ assigned grade. Each barplot shows the AUROC at which the status of a certain marker can be predicted from histology images using only histology grade.

An interesting observation from our stratification analysis is that the performance of ER predictor significantly increases from AUROC of 0.90 to 0.95 on low-grade cancer cases in the ABCTB cohort. This improvement could be attributed to a shift in the association structure between histological grade and ER status in both the cross-validation (TCGA-BRCA) and independent validation (ABCTB) cohorts (see **Figure 5B**). From the figure, the mutual-exclusivity association between ER status and histology grade is relatively stronger in the ABCTB cohort compared to the TCGA-BRCA cohort. This suggests that the model depends upon features associated with grade beyond those solely linked to PR status variations for prediction, potentially improving the ER model’s accuracy on cases in the ABCTB cohort. We observe a similar trend for our HER2 predictor. In the TCGA cohort, our HER2 status predictor was relatively less accurate with an average AUROC of 0.67, however, it became a relatively strong predictor with an AUROC of 0.77 on patients in the ABCTB cohort. Nonetheless, despite the improved predictive performance of the HER2 predictor on patients in the ABCTB cohort, the model’s accuracy remains poor for low-grade tumours, yielding an AUROC of 0.59.

### 2.7 TP53 mutation prediction in breast tumours is confounded by histology grade

We analysed the influence of histological grade on the TP53 mutation predictor by comparing the model performance over test cohorts stratified based on patients’ histological grade. TP53 predictor achieved state-of-the-art performance with an average AUROC of 0.81 and a narrow confidence interval (0.81-0.82). This demonstrates the TP53 predictor’s effectiveness in distinguishing mutant cases from wild-type cases from WSIs. This figure, while initially promising, is nuanced by a stratification analysis by histological grade. The model performance significantly declines to AUROC values of 0.73, 0.73 and 0.72, respectively for low, medium and high-grade tumours (FDR-corrected p-values of permutation testing p≪0.05). Such a drop, reminiscent of Simpson’s Paradox, indicates that the model relies on grade-associated features beyond those associated solely with TP53 mutational status variations for its prediction. Additionally, this also suggests that within each subgroup of patients, the model’s prediction is not more informative than using pathologist-assigned grade information for prediction, which yields an AUROC of 0.75 for predicting mutation status by itself (see **Figure 6**).

### 2.8 Endometrial cancer TP53 and PTEN mutation predictors are confounded by histology grade

We analysed the influence of histological grade on TP53 and PTEN mutation predictors by comparing the performance of these models on test cohorts stratified based on patients’ histological grade. These models predict TP53 and PTEN mutation status of patients in the TCGA-UCEC cohort with higher AUROC values of 0.788 (0.782-0.795) and 0.803 (0.798-0.808), respectively. Additionally, on the external validation cohort, these predictors achieve AUROC values of 0.70 (0.679-0.724) and 0.731(0.703-0.76), respectively. These results highlight the effectiveness of these predictors in inferring mutations in these genes from WSIs. However, after stratification analysis, the performance of both TP53 and PTEN predictors drops significantly in low and medium-grade tumours in the TCGA-UCEC cohort to AUROC values of (0.45, 0.51), (0.63,0.57), respectively in low and medium grade tumours (see **Figure 5A**). Likewise, in the CPTAC-UCEC cohort, we observe a drop in the predictive performance of these models.

An important observation is that the magnitude of the performance drop for these models varies across both cohorts. For example, the performance of the TP53 predictor drops on high-grade tumour cases in both the TCGA and CPTAC cohorts, however, in the CPTAC cohort we see a massive drop in performance (AUROC: 0.36). Similarly, the performance of the PTEN mutation predictor drops on medium-grade cases in both TCGA-UCEC and CPTAC-UCEC cohorts, however, in the CPTAC cohort we see a massive drop in performance (AUROC of 0.25). This massive drop in performance is likely to stem from the change in association structure between grade and TP53 and PTEN mutation status across cross-validation (TCGA-UCEC) and independent validation (TCGA-UCEC) cohorts as can be seen in **Figure 5B**. From the figure, in the TCGA-UCEC cohort low-grade cancer cases are more likely to have TP53 and PTEN mutations however we see roughly opposite trends for patients in the CPTAC-UCEC cohort where low-grade tumour cases are less likely to have TP53 and PTEN mutations.

### 2.9 The added predictive power of tumour receptor status predictors beyond pathologist grade assignments

Our analysis shows that the ER model’s predictions in low-grade tumours and the PR model’s predictions in low and medium-grade tumours are not substantially more informative than the pathologists’ grade assignments alone. To quantify this, we have established a baseline by predicting ER and PR status using only tumour grade as an input, which yields an AUROC benchmark of (ER: 0.76, PR: 0.70) and (ER: 0.79, PR: 0.71) respectively for patients in TCGA-BRCA and ABCTB cohorts (see **Figure 6**). Given that the baseline AUROC based on histology grade alone approaches that of our more complex model (ER: 0.87, PR: 0.79), it implies that benchmark of (ER: 0.76, PR: 0.70) and (ER: 0.79, PR: 0.71) respectively for patients in TCGA-BRCA and ABCTB cohorts (see **Figure 6**). Given that the baseline AUROC based on histology grade alone approaches that of our more complex model (ER: 0.87, PR: 0.79), it implies that the model’s intricate algorithms offer limited additional predictive value over traditional pathologist review. Additionally, this analysis suggests that a simple model that relies only on a patient’s histological grade can achieve high prediction accuracy without using a sophisticated machine learning algorithm.

### 2.10 Predictability of gene mutations based on pathologist-assigned grade

We analysed the predictability of gene mutation prediction based on the pathologist’s assigned grade. From our analysis, we found that histological grade shows significant predictability in predicting gene mutations in breast, lung adenocarcinoma and colorectal tumours as shown in **Figure 6**. For instance, TP53 mutations in breast tumours can be predicted with an AUROC of 0.75 which is close to AUROC of 0.81 obtained using a weakly supervised ML model. As the baseline AUROC derived solely from histology grade approaches that of our more complex model, it implies that the model’s intricate algorithms offer limited additional predictive value over traditional pathologist review. Similarly, TP53 and PTEN mutation in endometrial cancer can be predicted with high accuracy solely using histological grade in both cross-validation (TCGA-UCEC) and independent validation (CPTAC-UCEC) cohorts (see **Figure 6**). This analysis shows that for some gene mutations (e.g., TP53, etc.) in breast and endometrial cancer specimens, a simple predictor that uses only pathologist-assigned grade can achieve high accuracy, without using a sophisticated deep learning-based model.

### 2.11 Colorectal cancer mutation predictors are influenced by the density of mutations in other genes

We analysed the influence of mutational density of other genes on WSI-based mutation predictors by comparing the performance of these models on test cohorts stratified based on patients’ mutational load. From our analysis, in the TCGA-CRC cohort, WSI-based models achieve an AUROC value of 0.774 (0.764-0.785) for predicting BRAF mutations and 0.717 (0.711-0.722) for predicting TP53 mutations. However, detailed analysis reveals a significant challenge: for cases with low density of mutations in genes other than BRAF (denoted as 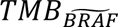), the BRAF predictor accuracy significantly drops to an AUROC of 0.65 (see **Figure 7A**). Similarly, the TP53 predictor behaves like a random predictor with an AUROC of 0.5 for cases with high TMB. In the CPTAC-CRC cohort, we observe similar trends, where the predictive performance of BRAF and TP53 predictors drops in cases with low and medium TMB, respectively. Apart from BRAF and TP53, the APC and KRAS mutation predictors are also influenced by TMB as can be seen in **Figure 7A**.

**Figure 7:**
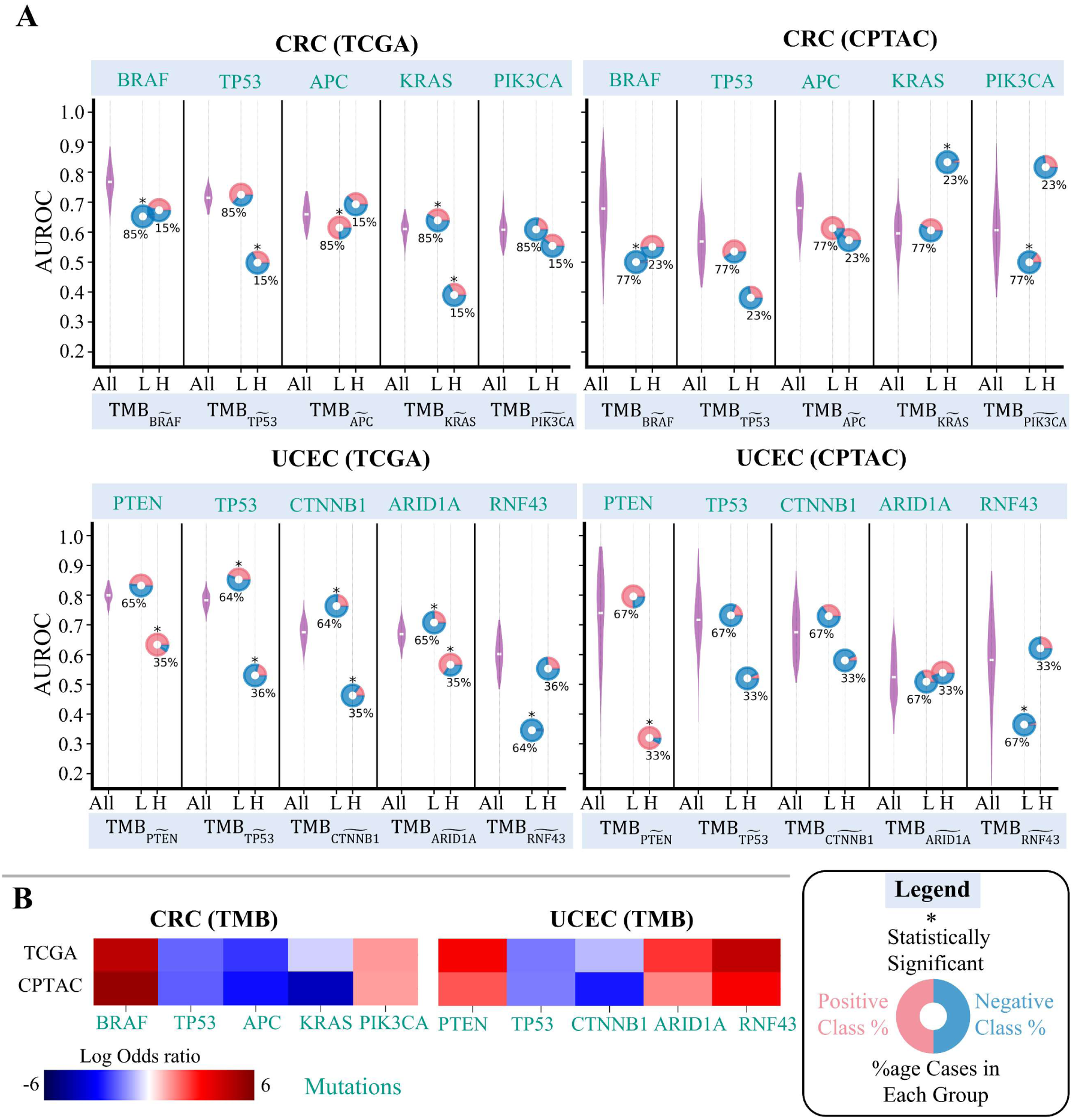
Plots illustrating the biased predictive performance of image-based biomarker predictors across patients with different tumour mutational burdens through stratification analysis. **A)** In the plots, AUROC values are illustrated on the y-axis, with the top x-axis indicating the prediction variables and the bottom x-axis showing patients’ stratification concerning tumour mutational burden. The predictive performance of each predictor on all the cases in the cohort (denoted by All in the plot) over 100 bootstrap runs is shown using a violin plot, whereas its performance in patients with high and low tumour mutational burden is depicted with a donut chart, with the centre representing the AUROC values. Donuts marked with * at the top indicate statistically significant results (FDR-corrected p-values of permutation testing *p* ≪ 0.05). Red and blue colours in each donut indicate the proportion of positive and negative cases in each stratified group based on prediction variables. **B)** Heatmaps highlighting the shift in the association structure between tumour mutational burden and gene mutations across two distinct datasets. Color intensity reflects the strength of association, with dark-red indicating strong co-occurrence and dark-blue indicating strong mutual exclusivity.

An interesting observation from our detailed analysis is that the performance of the KRAS predictor significantly increases from an AUROC of 0.63 to 0.83 on higher TMB cases in the CPTAC-CRC cohort. This improvement can be attributed to a shift in the association structure between TMB and KRAS mutation status in both TCGA-CRC and CPTAC-CRC cohorts (see **Figure 7B**). From the figure, the mutual-exclusivity association between KRAS mutation status and TMB is relatively stronger in the CPTAC-CRC cohort compared to the TCGA-CRC cohort. This suggests that the model’s predictions are influenced not just by KRAS mutation which is our target prediction variable but also by the overall mutation burden.

### 2.12 Endometrial cancer mutation predictors are influenced by the density of mutations in other genes

PTEN mutation predictor achieved higher AUROC values of 0.803 (0.798-0.808) and 0.731(0.703-0.76) in predicting the mutational status of patients from histology images in TCGA-UCEC and CPTAC-UCEC cohort, respectively. However, the predictor performance drops significantly for cases with low TMB in both TCGA-UCEC and CPTAC-UCEC cohorts to an AUROC of 0.63 and 0.32, respectively. Similarly, the performance of the TP53 predictor drops for cases with low TMB in both the TCGA-UCEC and CTPAC-UCEC cohort as can be seen in **Figure 7**. Apart from PTEN and TP53, the predictors for several other genes mutation predictors show varying accuracy across low and high TMB cases (see **Figure 7**).

## 3 Discussion

Through statistical analysis, we have demonstrated substantial interdependence among biomarkers as well as gene mutation statuses in breast, colorectal, endometrial, and lung adenocarcinoma samples, spanning across four cohorts: TCGA, METABRIC, MSK, and DFCI. We showed that the existence of such interdependencies in training data hinders the ability of current weakly supervised algorithms to accurately predict individual biomarker or gene mutation level genotype-to-phenotype mappings independent of other biomarkers. Consequently, biomarkers or gene mutation predictors based on histology images rely on the inherent association among the status of biomarkers on gene mutation in the training data. Predictors trained on such data are likely to capture a combined effect of multiple gene mutations of biomarkers that show mutual exclusivity or co-occurrence of molecular alterations in the training dataset. This imposes a strong prior on these predictors, assuming that associations present in training data will also persist for test samples. If that is not the case, these models may exhibit biased performance across sample populations with varying molecular characteristics. Through stratification analysis, we have demonstrated that the performance of several predictors significantly declines after grouping patients based on the status of a co-dependent marker and then analysing the model’s performance in each group (see **Figure 4** and **Figure S3**). Given the multi-faceted influence of molecular characteristics and other factors on the phenotypes observed in WSIs of cancer patients, it can be challenging for deep learning algorithms to discern individual biomarker-level genotype-to-phenotype mapping without appropriate causal adjustments.

Histology image-based biomarker predictors in breast tumours exploit histology grade-associated features for predicting the status of hormone receptors (ER and PR), rather than learning visual patterns solely linked to receptor status variation. Similarly, breast tumours’ TP53 mutation predictor and PTEN and TP53 mutation predictors for endometrial cancers are also confounded by histological grade. Results from other predictive models reveal similar trends (see **Figure 5**). This observation is not unique to our receptor status or mutation status prediction model but reflects a broader issue in the evaluation of machine learning models in computational pathology. Machine learning models risk capturing histological patterns associated with confounding variables (such as tumour grade), inadvertently conflating them with biomarkers of interest (e.g., ER, PR, TP53 and PTEN status). Such confounding factors resulting from causal or non-causal associations between prediction targets and other variables can obscure true predictive relationships, limit the generalisability of models, and inadvertently introduce bias. This underscores the importance of performing stratification analyses to ensure that the resulting models are equally generalisable across patient subgroups prior to deployment in clinical settings.

Our results show that models designed to predict gene mutations from histology images are susceptible to the influence of overall TMB or mutational density in other genes. For instance, we showed that the effectiveness of TP53 mutation predictors in colorectal tumours diminishes (AUROC of 0.5) in cases with high TMB (see **Figure 7**). This significant performance drop points to a confounding effect: the model’s predictions are influenced not just by TP53 mutation which is our target prediction variable but also by the overall mutation burden, i.e., mutations in other and even unrelated genes. Essentially, the predictor appears to use the broader mutational context or other factors as an indirect indicator of TP53 status, conflating TP53 mutations with general tumour characteristics linked to a low mutation load and its phenotypic effects on the tissue observable in the histology image. It is crucial to recognise that this phenomenon where predictive accuracy is influenced by the overall mutation burden is not unique to our TP53 prediction model. This effect, stemming from the dataset’s inherent correlation and inter-dependence among mutations, extends to other machine learning models (e.g., those predicting the status of mutations in BRAF, KRAS, PTEN etc) trained in a similar way. It is important to recognise such biases in predictive models and overcome them prior to the deployment of such models.

While external validation often serves as a benchmark for evaluating the performance of histology image-based biomarker predictors, it should not be the sole mechanism of validation and verification of WSI-based predictors of molecular status. For example, we demonstrated that our predictors for ER and PR status achieve state-of-the-art performance in both cross-validation (TCGA-BRCA) and a larger independent validation (ABCTB) cohort. Notably, these predictors showed slightly enhanced performance in the independent validation cohort (see **Figure 3** and **Figure 5**). However, upon closer examination, we found that this improvement stemmed from the stronger association between histology grade and ER status in the independent validation cohort compared to the cross-validation cohort. This underscores a limitation in the generalisability of histology image-based biomarker predictors: if the association structure between histology grade and receptor status weakens in another target dataset or sample population, the performance of these models may become erratic. This also suggests that the predictions of a sophisticated deep learning model may not be substantially more informative than a baseline predictor based on pathologist-assigned histological grade.

Achieving the goal of relying on routine WSI-based prediction of biomarkers without the need for more direct approaches (such as sequencing or immunohistochemistry) requires the development of significantly more sophisticated prediction algorithms. These methods should go beyond the mere correlation analysis and delve deeper into comprehending the causal structure of the problem. To achieve this, further research is imperative to explore innovative approaches capable of not only discerning associations but also uncovering causal linkages among various phenotypic patterns observed in WSIs and genetic alteration data. Additionally, these methods should address modelling associations in the label space, mitigating the impact of potential confounders that could lead to biased models.

Finally, in the context of deep learning models for the prediction of clinically actionable biomarkers from histology images, it is imperative to specify conceivable priors on the model stemming from the data. We emphasise that suitable stratification analyses are performed to ensure that the WSI-based predictors of molecular biomarkers are equally generalisable across patient subgroups before they can be deployed in clinical settings. Clinically, failure to account for interdependencies may lead to suboptimal decisions, necessitating appropriate usage guidelines for predictive models.

Our findings suggest revisiting model training protocols to recognize and adjust for biomarker interdependencies at all stages of model development—from problem definition to usage guidelines. This involves selecting diverse datasets to reflect clinical heterogeneity, defining prediction variables, or grouping patients based on significant co-dependencies, designing models to disentangle complex relationships, and implementing stringent stratification testing. To address these challenges, several strategies can be implemented.

Firstly, enhancing dataset diversity is crucial. This can be achieved by including samples from various demographics, disease stages, and treatment backgrounds, ensuring the training data encompasses the full spectrum of clinical scenarios while considering any differences in treatments or other variables that can bias the prediction model. Secondly, adopting multi-task and multi-label learning frameworks can help models learn shared representations while differentiating between correlated biomarkers, improving their ability to disentangle complex relationships. The machine learning model should report the full range of prediction variables rather than focusing on a single variable. This comprehensive reporting allows for a more nuanced understanding of the complex interdependencies among biomarkers and gene mutations, enhancing the model’s utility in clinical decision-making.

Thirdly, incorporating causal inference techniques into model development can provide a more robust understanding of the underlying causal structures, reducing the risk of confounding influences. Another approach is to group prediction variables that are causally dependent into higher-level patient groups or topics that are semantically or pathologically meaningful while ensuring an appropriate level of granularity in prediction. Additionally, clustering patients based on multi-biomarker profiles and predicting these clusters can ensure that patient groups are clinically relevant. Fourthly, rigorous stratification testing should be performed to evaluate model performance across different subgroups, identifying and mitigating biases before clinical deployment.

Finally, developing comprehensive usage guidelines and decision-support systems that account for potential interdependencies and model limitations will ensure that clinical decisions are informed by reliable and accurate predictions. By integrating these strategies, we can advance the development of predictive models that are both accurate and generalizable, ultimately improving patient care.

## 4 Methods

### 4.1 Patient Cohorts

We analysed data of four cancer types (breast, colorectal, lung adenocarcinoma and endometrial cancer), sourced from six cohorts: TCGA^40^, METABRIC^30,31^, COAD-DFCI^34^, MSK-LUAD^32^, CPTAC and ABCTB. Biomarkers and genes mutation status information, except for the ABCTB cohort, were collected from cBioportal [43], [44]. WSIs of Formalin-Fixed Paraffin-Embedded (FFPE) Haematoxylin and Eosin (H&E) stained tissue for TCGA atlas cases were collected from TCGA^40,41^, while for CPTAC atlas cases, they were retrieved from The Cancer Imaging Archive (TCIA)^17^. Within the ABCTB cohort, WSIs and receptor status (ER, PR and HER2 status) information were available for 2,303 patients. In terms of biomarkers, for breast tumours, ER, PR and HER2 status were recorded. For colorectal cases, MSI, hypermutation (HM), chromosomal instability (CIN), and CpG island methylator phenotype pathway (CIMP) activity statuses were documented. TMB information was available for all cases across all cancer types and cohorts.

### 4.2 Inter-gene mutational dependency analysis

We analysed the inter-dependency between the mutational status of genes using log2 odd ratio. Given the status of two biomarkers A and B, we calculated the log odd ratio (LOR) as follows:

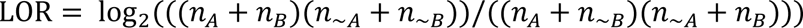

In the above equation, *n*_A_ and *n*_B_ indicates the number of cases with mutations in biomarkers *A* and *B*, respectively, while *n*_∼A_and *n*_∼B_ indicates the number of cases in the dataset where biomarkers A and B are not mutated. A higher positive LOR between gene pairs indicates mutation co-occurrence (i.e., if one gene is mutated the other is likely to be mutated), while a negative value signifies mutual exclusivity of mutation (i.e., if one gene is mutated, the other is less likely to be mutated).

In addition to the LOR analysis, we statistically assessed the inter-dependence among the mutational status of different genes using a two-sided Fisher exact test. All gene pairs were enumerated, and a Fisher exact test was performed on each pair. Subsequently, we reported the multi-hypothesis corrected p-values for each pair using the Benjamini/Hochberg method, with a significance threshold set at *p* ≪ 0.05.

### 4.3 Prediction of biomarkers and gene alteration from WSI

We assessed the predictability of biomarkers and gene alteration status from WSIs within their respective cohorts using two different algorithms with different principles of operation: CLAM^35^ and *SlideGraph*^∞36^. In order to avoid drawing conclusions specific to a certain approach or type of features, the predictive performance of both algorithms was evaluated over different types of features: deep features (convolutional neural network based encoder trained on ImageNet)^42^ and self-supervised features (a transformer-based model trained on histology images in a self-supervised manner)^43^. Our predictive pipeline comprises three main steps: 1) pre-processing of WSIs, 2) embedding of WSI patches, 3) biomarkers and gene mutation prediction from WSI using CLAM and *SlideGraph*^∞^.

#### 4.3.1 Preprocessing

In our pre-processing pipeline, utilising a U-Net-based segmentation model from TIAToolbox^44^, we first segment viable tissue areas of each WSI and exclude regions with artefacts (pen-marking, tissue folding, etc.). The model-generated tissue mask highlights informative tissue areas within the WSI using a pixel value of 1, while those with a value of 0 represent background or regions with artefacts. By leveraging these tissue masks, from each WSI we exact patches of size 512 × 512 pixels and 1,024 × 1,024 pixels at a spatial resolution of 0.50 microns-per-pixel (MPP). We selectively keep patches (both benign and tumour) that have over 40% viable tissue in terms of pixel proportion.

#### 4.3.2 Feature representation

We utilised various encoders to extract feature representation from WSI patches. Specifically, we employed ShuffleNet^45^, pretrained on ImageNet^42^ as a patch-level encoder to extract 1024-dimensional feature representation (deep features) from WSI patches of size 512 × 512 pixels. Additionally, we also extracted 768-dimensional self-supervised feature representation from each patch of size 1,024 × 1,024 pixels using CTransPath (a transformer-based self-supervised model trained on histology images)^43^.

#### 4.3.3 Predictive models

We trained *SlideGraph*^∞^and CLAM for predicting the status of different clinical biomarkers using both deep features and self-supervised features. In case of *SlideGraph*^∞^, we first construct a graph representation of the WSI, and then pass the WSI-graph to a Graph Neural Network for predicting the status of a certain biomarker as output. In cases where patients had multiple WSIs, we constructed a serial graph incorporating all WSIs and predicted the target label accordingly. In the case of CLAM, we bag all the WSIs belonging to the same patient and then predict the target label from the WSI-bag.

### 4.4 Training and validation of Image-based predictors

We trained and evaluated the performance of both *SlideGraph*^∞^and CLAM using 4-fold cross-validation, in which the dataset is partitioned into four 75/25 non-overlapping splits. In each cross-validation run the model is trained on 75% data, and the performance of the trained model is then assessed on the 25% test set. From the training dataset, we randomly select 10% of the data for validation. We trained the model for 300 epochs on the training set, with a batch size of 8 and a learning rate of 0.001 using the adaptive momentum-based optimiser ^46^. To limit overfitting, we stop the model training if its performance on the validation cohort is not improving over 10 consecutive epochs. We quantitatively assess the model performance on the test set using AUROC as a performance metric. We used the same train, validation, and test splits for both *SlideGraph*^∞^and CLAM.

### 4.5 Baseline predictors based on histology grade

To assess the predictability of biomarkers and gene mutation status based on histology grade, we employed a linear model (specifically, a support vector machine). This model utilises one-hot encoded histological grade as input to predict the status of certain a clinical biomarker as the target. We followed the same training and evaluation protocols used for our weakly supervised models. We quantify the model’s predictive performance using AUROC as a performance metric.

### 4.6 Stratification analysis to evaluate the impact of confounding factors

In predictive modelling, confounding variables can lead to false associations between the features and the target variable. Current weakly-supervised methods^6,7,47^ in computational pathology often overlook gene co-dependencies, focusing instead on predicting single-gene phenotypic effects. However, co-dependent genes’ phenotypic influence on histology images may confound associations between image features and target biomarker status, potentially resulting in a model that predicts the combined effect of multiple genes rather than a singular gene. These co-dependencies could also lead to models with poor generalisation performance, as the association structure among biomarkers may vary across datasets. To illustrate this, we analysed the performance of a machine learning model for a certain biomarker (e.g., ER status) concerning a certain stratification variable which we intend to test (e.g., TP53) using the algorithm provided in **Table 1**. For example, for ER predictor with TP53 as a stratification variable, we evaluated if the predictive performance of ER predictor varies with respect to TP53 status using permutation analysis. Specifically, we compare the observed AUROC values for the ER predictor in TP53 mutant and wild-type cases to the null distribution (obtained by permuting the dataset across 10,000 trials). If the model (e.g., ER predictor) varies significantly (multiple hypothesis corrected *p* ≪ 0.05) across patients’ subgroups (e.g., TP53 mutant and wild-type cases), then it indicates possible confounding effects or Simpson’s Paradox ^48,49^. We hypothesise that if confounding variables (i.e., the status of other biomarkers or clinicopathological parameters) do not influence the predictor, then the model’s accuracy within each stratified group should remain within the statistical variation of the overall data’s performance. However, if the model performance within each subgroup is substantially and statistically significantly different from the overall, then this indicates a possible confounding effect ^50,51^.

**Table 1:**
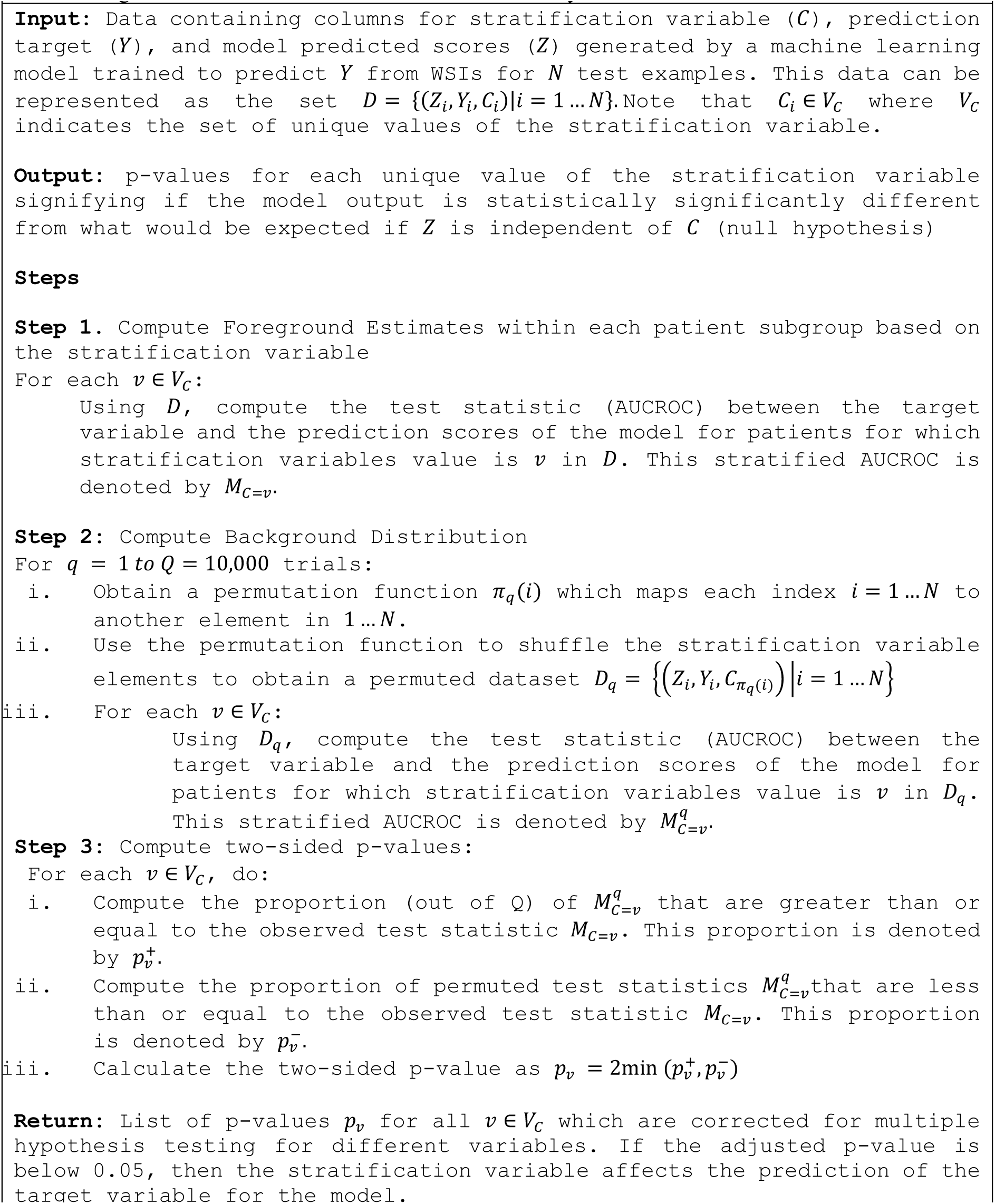
Algorithm of Permutation Test for Stratification Analysis.

We examined three key factors that could introduce bias into an ML model. First, the bias due to interdependency among biomarkers and somatic mutation status of genes in the training dataset. Second, a likely bias due to patients’ tumour histological grades. Third, an expected bias due to the TMB of a cancer patient. To assess the influence of interdependence among biomarker statuses on model predictive performance, we select the model with the highest AUROC score for each biomarker and run a permutation test, treating other biomarkers with co-dependent statuses as confounding variables. Subsequently, to analyse the influence of histological grade on histology image-based biomarker predictors, we employ a similar approach, utilising histology grade as a confounding variable. Finally, to evaluate the impact of TMB on histology image-based biomarker predictors, we first calculate patient-level TMB excluding genetic alterations of the gene of interest used for prediction, then use this 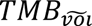 as a confounding variable. Based on 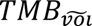 we divide the patients into low and high TMB cases using a threshold of ten mutations per megabase.

## Acknowledgements

MD would like to acknowledge the PhD studentship support from GlaxoSmithKline (GSK) and the Department of Computer Science at the University of Warwick. FM and NR were partially supported by the PathLAKE digital pathology consortium which was funded from the Data to Early Diagnosis and Precision Medicine strand of the government’s Industrial Strategy Challenge Fund, managed and delivered by UK Research and Innovation (UKRI). FM also acknowledges funding support from EPSRC EP/W02909X/1.

## Authors’ contributions

FM and MD designed the study with support from co-authors. FM and MD wrote the code and analysed the experimental results. NR, KB and FM secured the funding. FM and NR supervised the study. ST provided pathological and oncological input. MD and FM visualised and verified the underlying data. MD and FM drafted the manuscript with input from co-authors. All authors had full access to all the data in the study and had the final decision to submit for publication.

## Inclusion and diversity

We support inclusive, diverse, and equitable conduct of research.

## Data and code availability

- Whole slide images (WSIs) of TCGA patients used in the study can be downloaded from the NIH Genomic Data Common Portal at this link: https://portal.gdc.cancer.gov/ with the manifest details included in the supplementary material.
- The genomic data and clinical data of patients in TCGA, METABRIC, COAD-DFCI, MSK-LUAD and CPTAC cohorts can be downloaded from cBioPortal https://www.cbioportal.org/. Patient IDs and field details are included in the supplementary materials.
- ABCTB data and images were obtained from the Australian Breast Cancer Tissue Bank. The corresponding author could be contacted to facilitate access to ABCTB data.
- All the experiments were conducted using Python using PyTorch Geometric library and TIAToolbox. Code and documentation of all Python scripts used in the study can be found at: https://github.com/engrodawood/HistBiases.
- Any additional information required to reproduce the data reported in this work is available from the corresponding author upon request.

