## Supplementary Materials for "Buyer Beware: confounding factors and biases abound when predicting omics-based biomarkers from histological images"

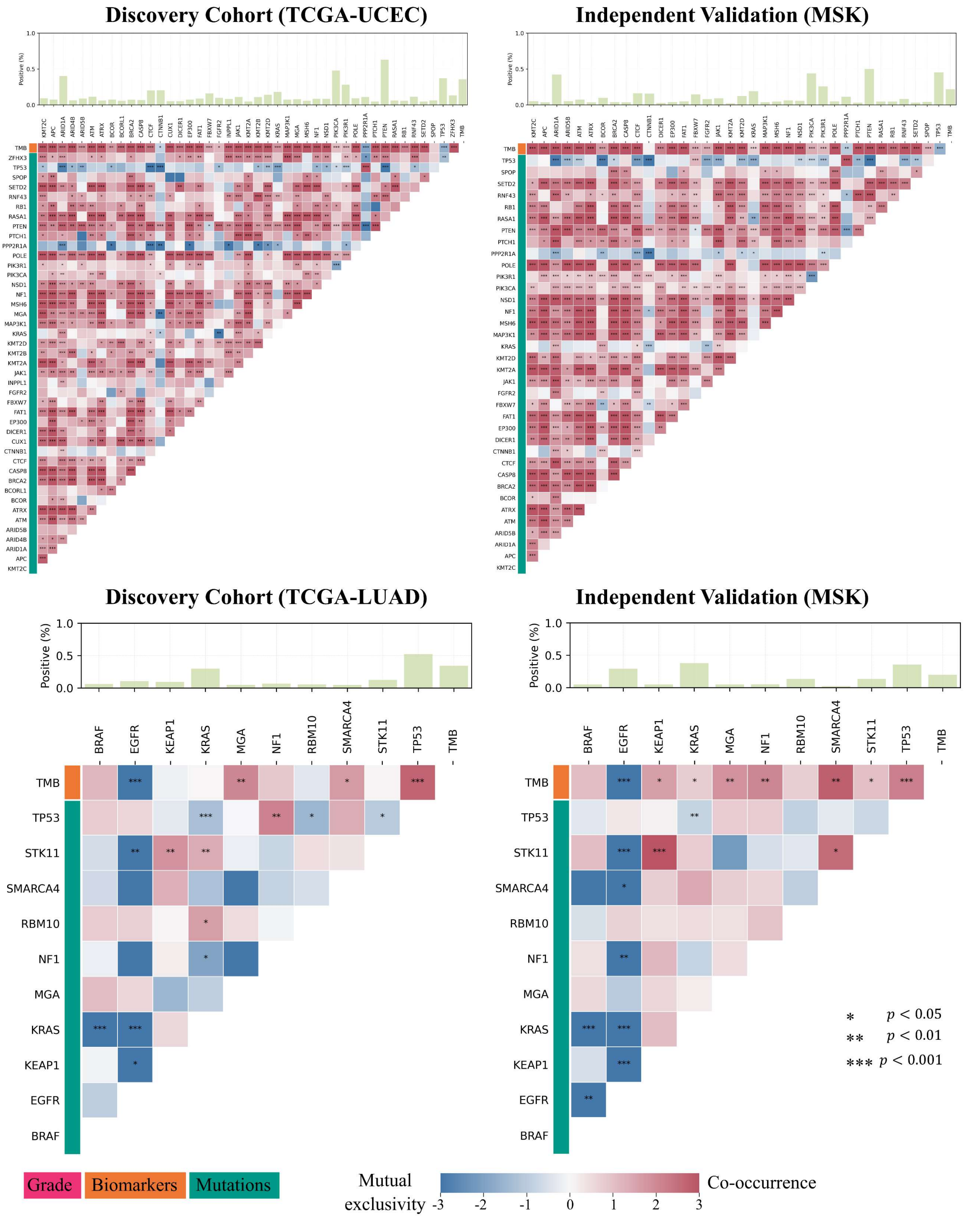


**Figure S1:** Association heatmaps of biomarkers and gene mutations status across tissue types and datasets, related to **Figure 2**.

The heatmaps display a set of genes along x and y axes. Cell colours within the heatmap illustrate the relationship and strength of association, with dark-red colours for co-occurrence and dark blue for mutual exclusivity. Heatmap cells marked with asterisks indicate significant associations (two-sided Fisher's exact test multiple hypothesis corrected p-value <0.05).

The top bar above each heatmap shows the percentage of cases mutated for a specific gene in case of gene mutations.

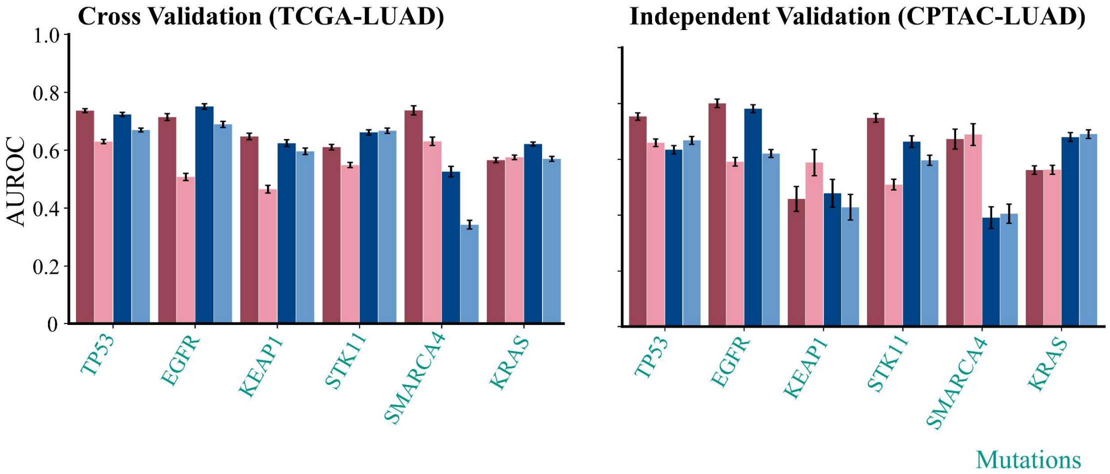


**Figure S2:** Quantitative results of prediction biomarkers/mutations from histology images related to **Figure 3**.

Each barplot shows the area under the receiver operating characteristic curve (AUROC) at which a certain clinical markers has been predicted using two weakly supervised model each with two different feature repsentation (i.e., using ShuffleNet pretrained on natural images as patch-level encoder, and CTransPath a transformer based model pre-trained on histology images using self-supervised learning).

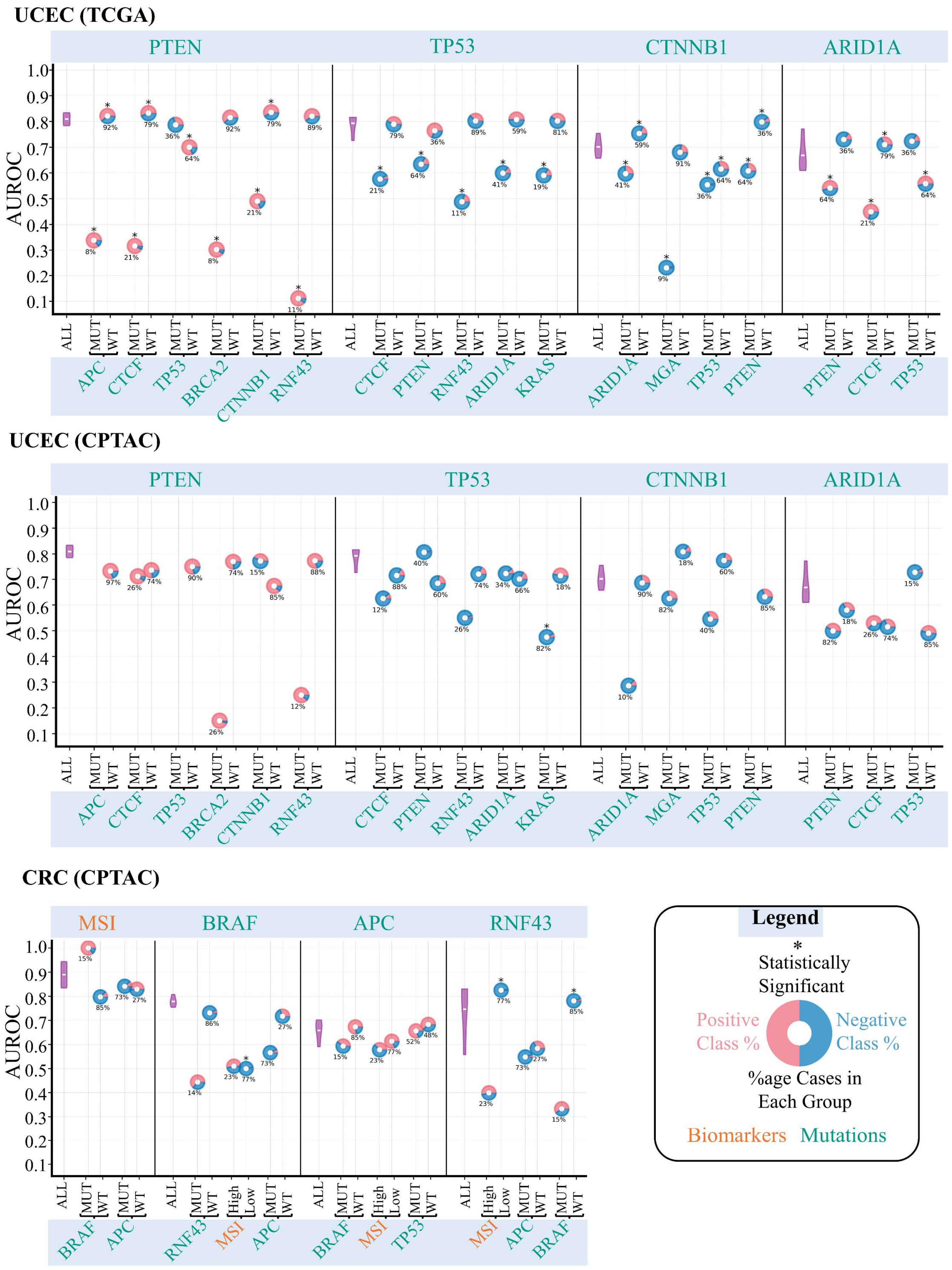


**Figure S3:** Plots showcasing stratification analysis of several histology image-based biomarker predictors concerning other interdependent biomarkers, related to **Figure 4**.

In the plots, AUROC values are illustrated on the y-axis, with the top x-axis indicating the prediction variables and the bottom x-axis showing the stratification variables. The predictive performance of each predictor on all the cases in the cohort (denoted by All in the plot) over 100 bootstrap runs is shown using a violin plot, whereas its performance in different stratified groups is depicted with a donut chart, with the centre representing the AUROC values. Donuts marked with * at the top indicate statistically significant results (FDR-corrected p-values of permutation testing << 0.05). The percentage values at the bottom of the donut indicate the proportion of positive (MUT/High) or negative (WT/Low) cases relative to the status of the stratification variables. Red and blue colours in each donut indicate the proportion of positive and negative cases in each stratified group concerning prediction variables.
